# Genome co-adaptation and the evolution of methicillin resistant *Staphylococcus aureus* (MRSA)

**DOI:** 10.1101/2025.07.21.665857

**Authors:** Seungwon Ko, Elizabeth A. Cummins, William Monteith, Samuel K. Sheppard

## Abstract

**Background:** Antimicrobial resistance (AMR) in bacterial pathogens is a major threat to global health, rendering standard treatments ineffective and increasing the risk of severe infection or death. Resistance is often conferred by genes that are transferred horizontally among species and strains. However, for many bacteria, little is known about the genetic variation that potentiates resistance gene acquisition and accommodates acquired genes in the coadapted recipient genome.

**Results:** Here we introduce a new bioinformatics genome-wide association study approach with Guided Omission of Linkage Disequilibrium (GOLD-GWAS) that masks covarying alleles explained by coinheritance and genome proximity to reveal genes where covarying sequence likely represents functional linkage between loci, consistent with epistasis. Analysing 806 *Staphylococcus aureus* isolate genomes, including methicillin-resistant (MRSA) and methicillin-susceptible (MSSA) strains, we identify genes that covary with the presence of the acquired staphylococcal cassette chromosome *mec* (SCC*mec*) that houses the *mecA* resistance gene.

**Conclusions:** Uncovering known and new gene-gene associations, we demonstrate how resistance can involve genetic coalitions beyond the well-known AMR genes. Understanding how genome change, here extrinsic resistance cassettes, are integrated within coadapted bacterial genomes is an important step towards mitigating AMR evolution by identifying novel genetic targets for risk prediction, diagnosis and therapy.

## Background

Bacterial populations can exhibit considerable trait variation. In some cases, closely related strains of the same species can vary from harmless commensals to important pathogens (1–3). Understanding how mutation and horizontal gene transfer (HGT) give rise to divergent phenotypes is a major focus in microbiology, with modern genomics linking gene function to trait variation in natural populations (4). Among the most problematic bacterial phenotypes is antimicrobial resistance (AMR), with projections that treatment failure will be associated with an estimated 50 million deaths worldwide by 2050 (5). Many of the genes, alleles, and polymorphisms underlying resistance are known. However, the function of genes depends on genomic context, and there is increasing evidence that the evolution of AMR may involve multiple genes, even for well characterized mechanisms (6–9).

In some well-characterized instances, a single gene or nucleotide polymorphism can increase resistance (10,11). Given this capacity for genomic plasticity, it can be tempting to view genes as modular elements that can be ‘plugged in’ or ‘switched on’ to confer resistance. However, this view oversimplifies the reality, as genes interact within genomes to confer phenotypes. This can be a basic additive effect where genes independently contribute to a phenotype, or non-additive effects where the effect of one gene or allele depends on another. Phenotypes conferred by non-additive gene effects can be either synergistic, where the sum of gene effects exceeds their individual contributions, or epistatic, where there is functionally interdependence and the action of one gene depends upon the other(s). Both can be associated with AMR (12–17).

Among the best-known AMR pathogens is methicillin-resistant *Staphylococcus aureus* (MRSA), a major cause of hospital acquired infections (18–20). The principal genetic driver of methicillin resistance is the *mecA* gene, which codes for a modified penicillin-binding protein (PBP2a) (21). This gene is commonly transported among *Staphylococcus* species in the staphylococcal cassette chromosome *mec* (SCC*mec*) (22–24), integrating into the recipient chromosome via sequence-specific recombination with *rlmH* (*orfX*) (25–27) (Supplementary Figure 1). The distribution of SCC*mec* in staphylococcal populations is a balance of forces favouring acquisition (28,29) and the fitness cost to the recipient strains (30–33). Understanding these opposing forces and the genetic consequences of SCC*mec* acquisition in natural populations requires large-scale comparative genomics.

Genome-wide association studies (GWAS) have been used to understand the genetics underlying phenotype variation in bacteria for over a decade (34). Bacterial GWAS has been applied to identify genetic determinants of AMR in pathogens, improving understanding of the emergence and spread of well-known resistance determinants, including SCC*mec* in staphylococci (35–41). While these studies may also highlight genes and alleles that covary with known AMR genes, this has seldom been an explicit aim. With increasing appreciation of the importance of gene-gene interactions in integrated bacterial genomes, there has been more emphasis upon understanding functional gene networks and identifying putative epistasis in population genomic datasets (42,43).

Genome-wide covariation analysis is conceptually simple. Essentially it involves, comparing genomes and identifying alleles that are found together, i.e. when ‘A’ is present at one locus ‘B’ is typically present at another. To make these findings biologically relevant it is important to compare the covariation signal to that which is expected by chance. Most co-variation is the result of co-inheritance or physical proximity on in the genome, not epistasis, and this can dominate signals in basic GWAS models. Here, we address this with a novel method that enhances traditional bacterial GWAS for co-variation analyses. Specifically, our approach (GOLD-GWAS) incorporates quantification of genome-wide linkage disequilibrium (LD) decay. That is to say, as the physical distance increases between two alleles they are less likely to have been coinherited because recombination will be more likely to have shuffled segments of DNA, which occurs by HGT in most bacteria. Having masked covariation resulting from LD, our approach identifies genes that covary with *SCCmec* in *S. aureus,* potentially indicative of potentiating or compensatory genome change. Using this approach, we are able to characterize the genomic landscape of AMR coadaptation, infer functional significant genes and identify putative epistatic loci.

## Results

### Testing genome masking with simulated data

To evaluate the performance of the Guided-Omission of LD GWAS (GOLD-GWAS) approach, we tested whether the method could detect an artificially generated covarying site within simulated bacterial genomes. We simulated a dataset comprising 1,000 genomes of 1 Mbp each, generated with a clonal genealogy under a coalescent model with recombination rate of R = 0.01 and a site-specific mutation rate of θ = 0.001 (44). An artificial site of covariation was constructed by identifying a polymorphism with minor allele frequence (MAF) > 0.2 within a 10 kbp range of the 100 kbp position. The polymorphism at 93,696 bp fulfilled these criteria and eleven artificial covarying sites between 600,000 bp and 600,100 bp at 10 bp intervals were created to covary with the 93,696 bp site. These artificial covarying sites were generated independently for each site with a 95% probability, to avoid numerical errors arising from perfect covariation. GOLD-GWAS was then applied to these simulated data with artificial covarying sites. From the GOLD-GWAS output, mapping k-mers to a reference genome demonstrated that our method effectively masked the target region with no k-mers mapping between 83,696 and 103,696 bp. Masking GWAS also identified accurately the artificial covarying site at 600,000 – 600,100 bp with 10 significantly associated k-mers present with -log(p-values) > 5.13 (Supplementary Figure 2).

### LD in *S. aureus* genomes declines to and equilibrates after 8790 bp

A total of 806 *S. aureus* whole genomes were chosen from the NCBI reference sequence database to represent both SCC*mec*-positive (n = 426) and SCC*mec*-negative isolates (n = 380). All assemblies were aligned to the NCTC 8325 reference genome before variant calling and 10% of all identified polymorphisms were randomly sampled for LD analysis. The average R^2^ value (a measure of LD) was determined for the data set, which fell to approximately 0.149 after 100,000 bp (Figure 1A). The gradient of the log-transformed graph of LD decay was calculated to be -0.0726 (Figure 1B). From these values, the overall LD range of *S. aureus* in our dataset was estimated to be 8790 bp, calculated as the intercept of the average R^2^ value and the fitted R^2^ logarithmic bp decay. Our estimate corresponds with a previous study of LD decay in *S. aureus* where no LD was reported between SNPs with distance greater than 10 kbp (45).

**Figure 1:**
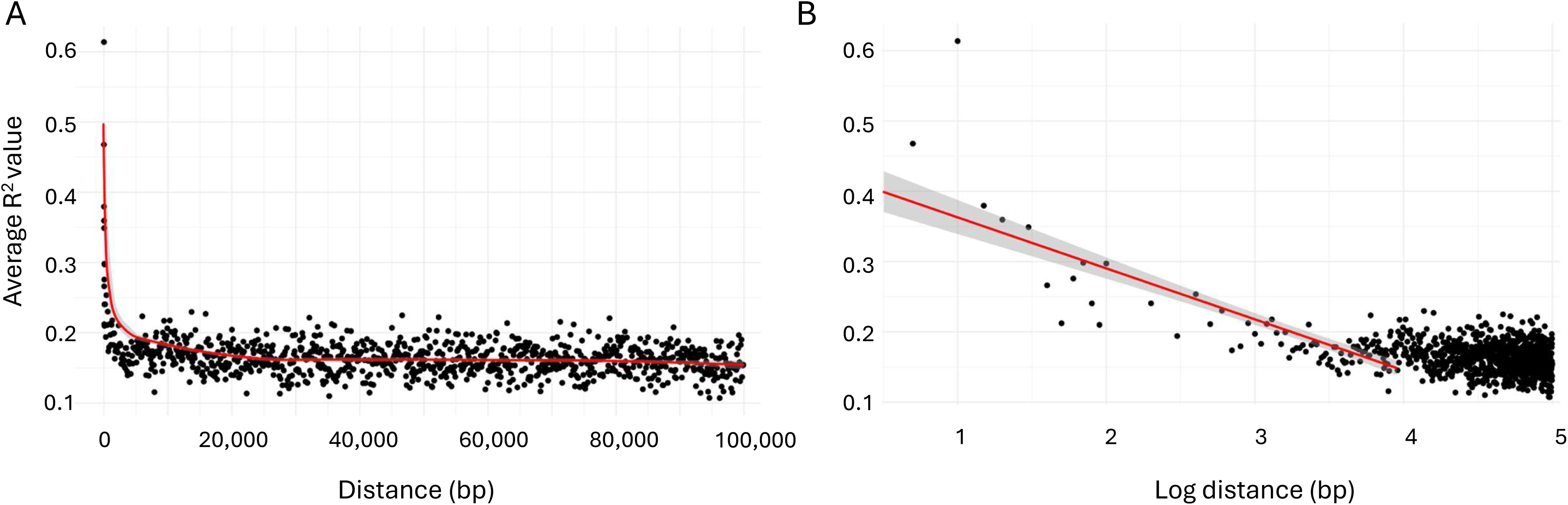
Distribution of linkage disequilibrium values (R^2^) across the genome as a function of the distance between single nucleotide polymorphism pairs in *S. aureus*. (A) Relationship between the linear distance between two SNPs and the average R^2^ value where each point is the R^2^ value for an individual distance category. Red line represents a polynomial trend line. (B) Relationship between the logarithm of the distance between two SNPs and the average R^2^ value. Red line represents the linear trend line fitted after excluding data points with R^2^ < 0.2.

### GOLD-GWAS identifies covariation associated with known epistatic sites in *S. aureus*

The GOLD-GWAS method was tested using biological data to identify genomic covariation among known epistatic sites. The sites selected for this test were: (i) *divIVA,* which play critical roles in cell division and polarity (46); (ii) *secA2,* which encodes an accessory secretion ATPase (SecA2) involved in the secretion of major autolysins such as p60 (CwhA) and MurA (NamA) both of which are key factors in bacterial cell wall remodelling (47) (48). The *divIVA* loci were chosen as the target region to evaluate whether the GOLD-GWAS pipeline could detect genomic co-variation within *secA2,* linked to *divIVA*.

Observed p-values from GOLD-GWAS closely followed expected values from a theoretical χ^2^-distribution up to approximately -log(p-value) = 2, beyond which points deviated from the null (Figure 2A). The sigmoidal distribution lacked clear ‘shelves’, confirming that there was no poorly controlled confounding population structure. Mapping k-mers to a reference genome (Figure 2B) revealed that k-mers significantly associated with *divIVA* carriage were found in the previously characterised epistatic region of *secA2* and notably *murC*, another gene involved in peptidoglycan synthesis (47). Removal of the *divIVA* locus and it’s LD regions ensured demonstrated that these genes emerged as independent associations - separate of the effects of LD. This is consistent with coadaptation with *divIVA* as previously described (46–48) and the utility of GOLD-GWAS to uncover biologically significant covariation in bacterial genomes. Furthermore, matching k-mers to the genes they are found in (Figure 2C) revealed that many of the top GOLD-GWAS hits had related functions, broadly linked to peptidoglycan biosynthesis (Table 1). Moreover, the genomic location of the top ranked genes (-log(p-value) > 200, beta > 0.5) indicated clearly that the significant associations with *divIVA* have not arisen solely from genomic proximity to the target region (Figure 2D).

**Figure 2:**
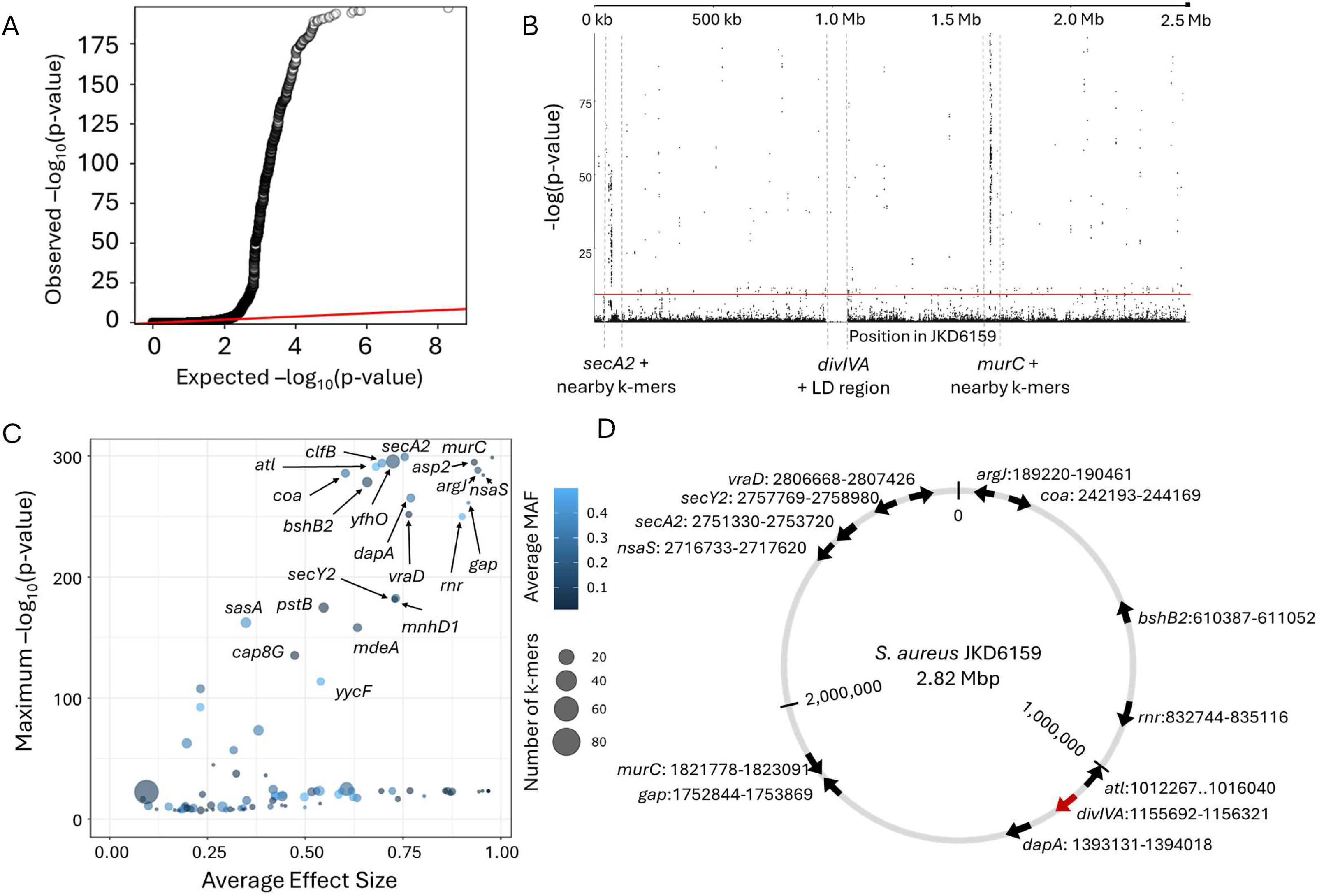
Summary of GOLD-GWAS after masking *divIVA* and associated LD regions. (A) Quantile-quantile plot comparing the expected -log(p-values) with observed -log(p- values) from GOLD-GWAS. The red diagonal line indicates where the expected and observed values are equal. (B) Manhattan plot demonstrating the statistical significance association for selected variants arranged in order on the reference genome JKD6159. Each dot represents a k-mer and the red line represents the threshold for significance. The positions of *murC*, *secA2*, *divIVA* and its LD region are indicated. (C) Plot of genes associated with *divIVA*. Minor allele frequencies (MAF) are shown by colour gradient and dot size represents the number of k-mers mapped to the gene. (D) Genomic location and orientation of genes covered by associated k-mers with a likelihood ratio test -log(p-value) > 200 and average effect size (beta) > 0.5 that exist within the *S. aureus* JKD6159 genome. Position of *divIVA* is indicated by the red arrow.

**Table 1:** Genes containing k-mers significantly associated with the presence of *divIVA* in *S. aureus* genomes (-log(p-value) > 200), ordered by descending maximum p-value.

| Gene | Description |
| --- | --- |
| *secA2* | Accessory Sec translocase SecA2 |
| *murC* | UDP-N-acetylmuramate-L-alanine ligase |
| *yfhO* | Lipoteichoic acid-specific glycosyltransferase YfhO |
| *asp2* | Accessory Sec system protein Asp2 |
| *clfB* | MSCRAMM adhesin clumping factor ClfB |
| *atl* | Bifunctional autolysin; cleaves peptidoglycan during cell division |
| *argJ* | Bifunctional glutamate N-acetyltransferase/amino acid acetyl-transferase |
| *coa* | Staphylocoagulase; cleaves fibrinogen |
| *nsaS* | Nisin susceptibility-associated sensor histidine kinase NsaS |
| *bshB2* | Bacillithiol biosynthesis deacetylase BshB2 |
| *dapA* | 4-hydroxy-tetrahydrodipicolinate synthase |
| *gap* | Type I glyceraldehyde-3-phosphate dehydrogenase |
| *vraD* | Peptide resistance ABC transporter ATPase subunit VraD |
| *rnr* | Ribonuclease R |

MSCRAMM: Microbial Surface Components Recognizing Adhesive Matrix Molecule.

### Masking SCC*mec* and the conserved LD region detects the SCC*mec* insertion site

In this study, the GOLD-GWAS pipeline begins with the computational masking of SCC*mec* and associated LD regions. LD regions are parameterised from the SCC*mec*-positive isolate genomes and observed p-values from GWAS were well controlled at low p-values (p < 1.5) consistent with the expected distribution under the null hypothesis. The even distribution of p-values suggested good control for potential confounding effects of population structure (Figure 3A). Mapping k-mers to the *S. aureus* SCC*mec*-positive reference genome showed that the masking effectively removed significant k-mers within the SCC*mec* and associated LD region (Figure 3Ci). A similar result was observed when k-mers were mapped to the SCC*mec*-negative reference genome apart from a single significant k-mer covering the attB site within *rlmH* (Figure 3Di). As the neighbouring sequences of *rlmH* are highly conserved, the computational masking covers the entire LD region surrounding the SCC*mec* region in both SCC*mec*-positive and SCC*mec*-negative isolates (49) (Figures 3Ci, Di). However, the attB site within *rlmH* is not conserved as it serves as the recombination site for SCC*mec* integration and therefore is present exclusively in SCC*mec*-negative isolates. Hence, the significant -log(p-value) of 9.5 observed for the k-mer that covers attB due to the strong negative correlation of this site with SCC*mec* carriage (Figure 3Di). This association effected the ranking of genes significantly associated with SCC*mec* carriage due to the extremely high (74.2) likelihood ratio test (LRT) -log(p-value) of *rlmH* that outranked covariation signals in other genes (Figure 3B). The genomic locations of the genes covered by significantly associated k-mers in both reference genomes demonstrated that these genes are distributed throughout the genome. The mean distance between genes was 187 kbp, with maximum and minimum distances of 782 kbp and 1.7 kbp respectively (standard deviation = 235 kbp). None of these genes were positioned within 10 kbp of *SCCmec* except for *rlmH*. Excluding the attB site in *rlmH,* this spatial separation indicated that significant hits are highly unlikely to result solely by physical proximity (Figures 3Cii, Dii).

**Figure 3:**
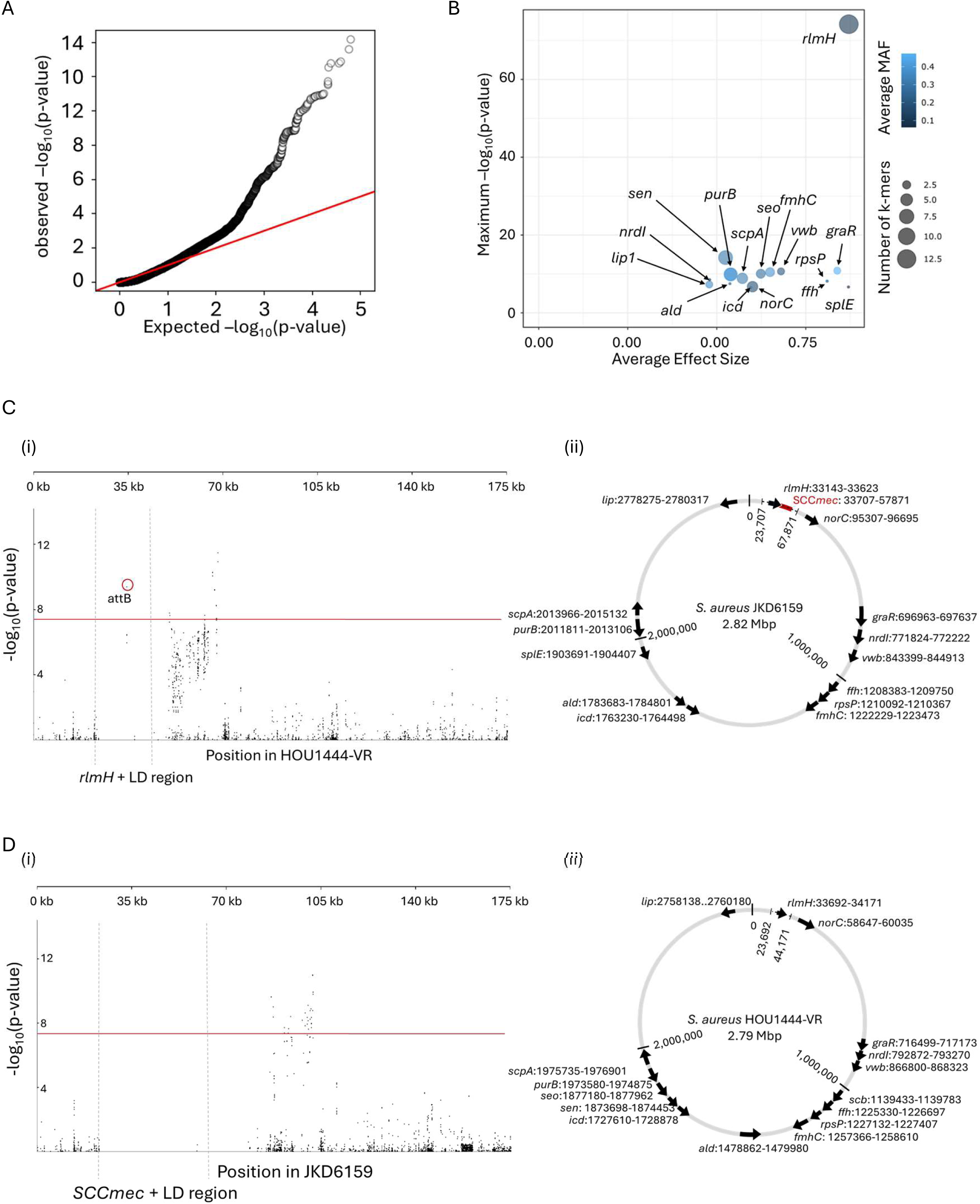
Summary of GOLD-GWAS results after masking SCC*mec* and conserved LD regions. (A) Quantile-quantile plot comparing the expected -log(p-values) with observed - log(p-values) from GOLD-GWAS. The red diagonal line indicates where the expected and observed values are equal. (B) Plot of genes associated with SCC*mec* carriage. Minor allele frequency (MAF) values are depicted by the colour gradient and dot size represent the number of k-mers mapped to the gene. (C) Manhattan plot and corresponding genomic map for the SCC*mec*-positive JKD6159 genome. (i) Manhattan plot demonstrating the statistical significance association for selected variants arranged in order on the 0–175 kbp region of the *S. aureus* JKD6159 genome. Th position of *SCCmec* and its associated LD regions are indicated. Each dot represents a k-mer and the red line represents the threshold for significance. (ii) Genomic location and orientation of significantly associated genes in SCC*mec*-positive JDK6159. Position of SCC*mec* is indicated by the red block and masked regions by dotted lines. (D) Manhattan plot and corresponding genomic map for the SCC*mec*-negative HOU1444-VR genome. (i) Manhattan plot demonstrating the statistical significance association for selected variants arranged in order on the 0–175 kbp region of the *S. aureus* HOU1444-VR genome. The position of *rlmH* and its associated LD region are indicated. The red circle highlights the k-mer identified as the attB site. (ii) Genomic location and orientation of significantly associated genes in SCC*mec*-negative HOU1444-VR. Masked regions are indicated by dotted lines.

### Masking SCC*mec* and *rlmH* detects genes that covary with SCC*mec* carriage

To improve the detection of sites that covary with SCC*mec,* beyond the attB site in *rlmH*, we masked SCC*mec* and *rlmH* alongside their respective LD regions using GOLD-GWAS. Again, observed p-values followed the expected distribution under the null hypothesis at low p-values (p < 1.5) with the even distribution confirming adequate control for population structure (Figure 4A). Mapping k-mers to the SCC*mec*-positive reference genome demonstrated effective removal of significant k-mers within SCC*mec* and associated LD regions (Figure 4Ci). Only 2 of the 172,224 total k-mers fell within the masked region with extremely low non-significant -log p-values of 0.04 and 0.33. Mapping k-mers to the SCC*mec*-negative reference genome also showed that no significantly covarying k-mers mapped to the *rlmH* gene and associated LD regions and only 3 of 172,224 k-mers were present with non-significant -log p-values of 0.24, 0.026, and 0.065 (Figure 4Di). The significantly covarying k-mers mapped to 16 genes (Table 2). The *sen* gene, which produces Staphylococcal enterotoxin type N, displayed the highest LRT value (-log(p-value) = 14.2, Figure 4B), while *splE*, which produces serine protease SplE, exhibited the largest effect size (beta = 0.87, Figure 4B). The chromosomal distribution of the 16 top-ranked genes in both SCC*mec*-positive and -negative reference genomes showed that the covarying genes were widely distributed across the genome. The mean distance between genes was 200 kbp, with maximum and minimum distances of 782 kbp and 1.7 kbp respectively (standard deviation = 239 kbp). No k-mers were located within 10 kbp of the insertion site by *rlmH* (Figures 4Cii, Dii).

**Figure 4:**
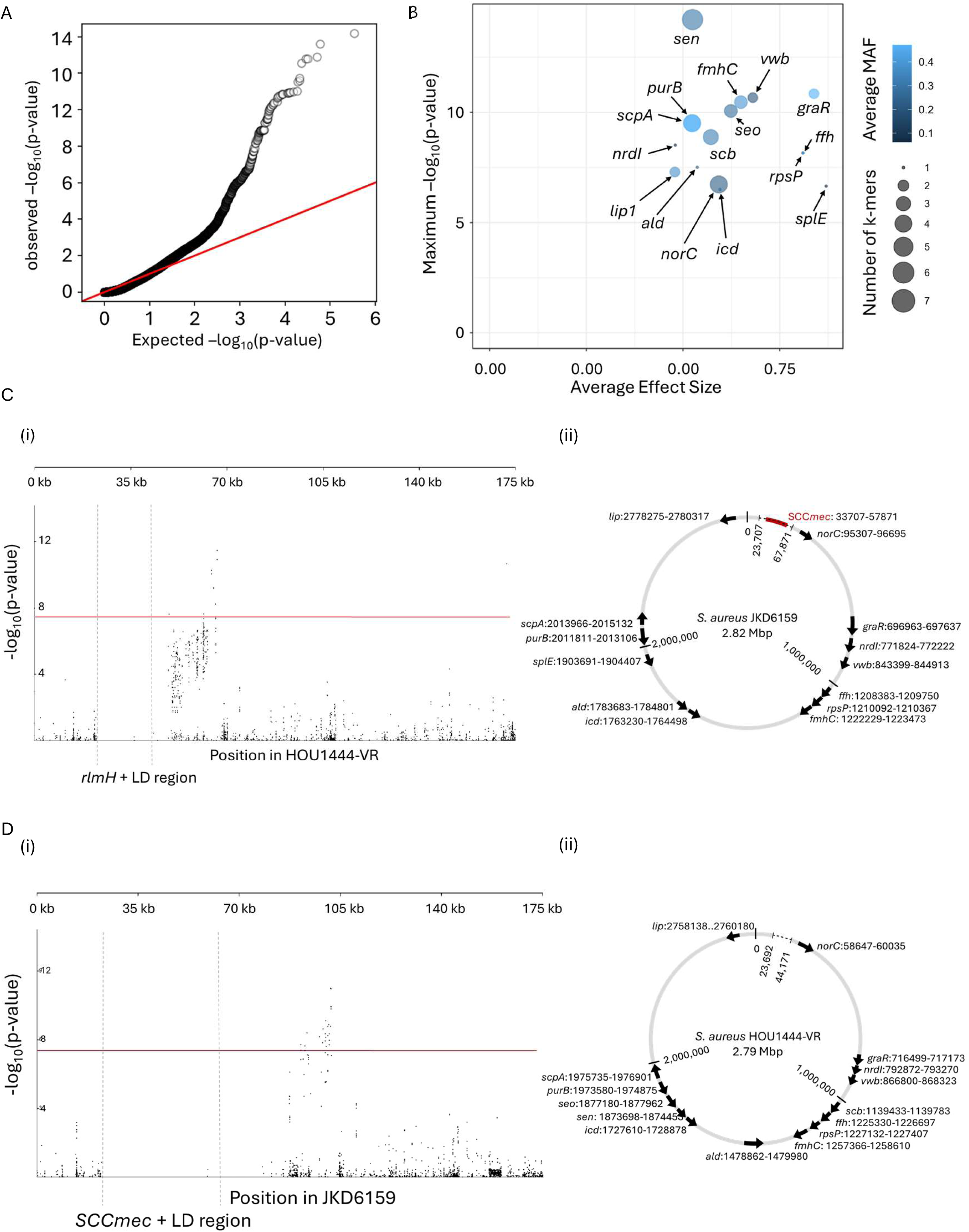
Summary of GOLD-GWAS results after masking SCC*mec, rlmH* and associated LD regions. (A) Quantile-quantile plot comparing the expected -log(p-values) with observed -log(p-values) from GOLD-GWAS. The red diagonal line indicates where the expected and observed values are equal. (B) Plot of genes associated with SCC*mec* carriage. Minor allele frequency (MAF) values are depicted by the colour gradient and dot size represent the number of k-mers mapped to the gene. (C) Manhattan plot and corresponding genomic map for the SCC*mec*-positive JKD6159 genome. (i) Manhattan plot demonstrating the statistical significance association for selected variants arranged in order on the 0–175 kbp region of the *S. aureus* JKD6159 genome. Th position of *SCCmec* and its associated LD regions are indicated. Each dot represents a k-mer and the red line represents the threshold for significance. (ii) Genomic location and orientation of significantly associated genes in SCC*mec*-positive JDK6159. Position of SCC*mec* is indicated by the red block and masked regions by dotted lines. (D) Manhattan plot and corresponding genomic map for the SCC*mec*-negative HOU1444-VR genome. (i) Manhattan plot demonstrating the statistical significance association for selected variants arranged in order on the 0–175 kbp region of the *S. aureus* HOU1444-VR genome. The position of *rlmH* and its associated LD region are indicated. (ii) Genomic location and orientation of significantly associated genes in SCC*mec*-negative HOU1444-VR. Masked regions are indicated by dotted lines.

**Table 2:**
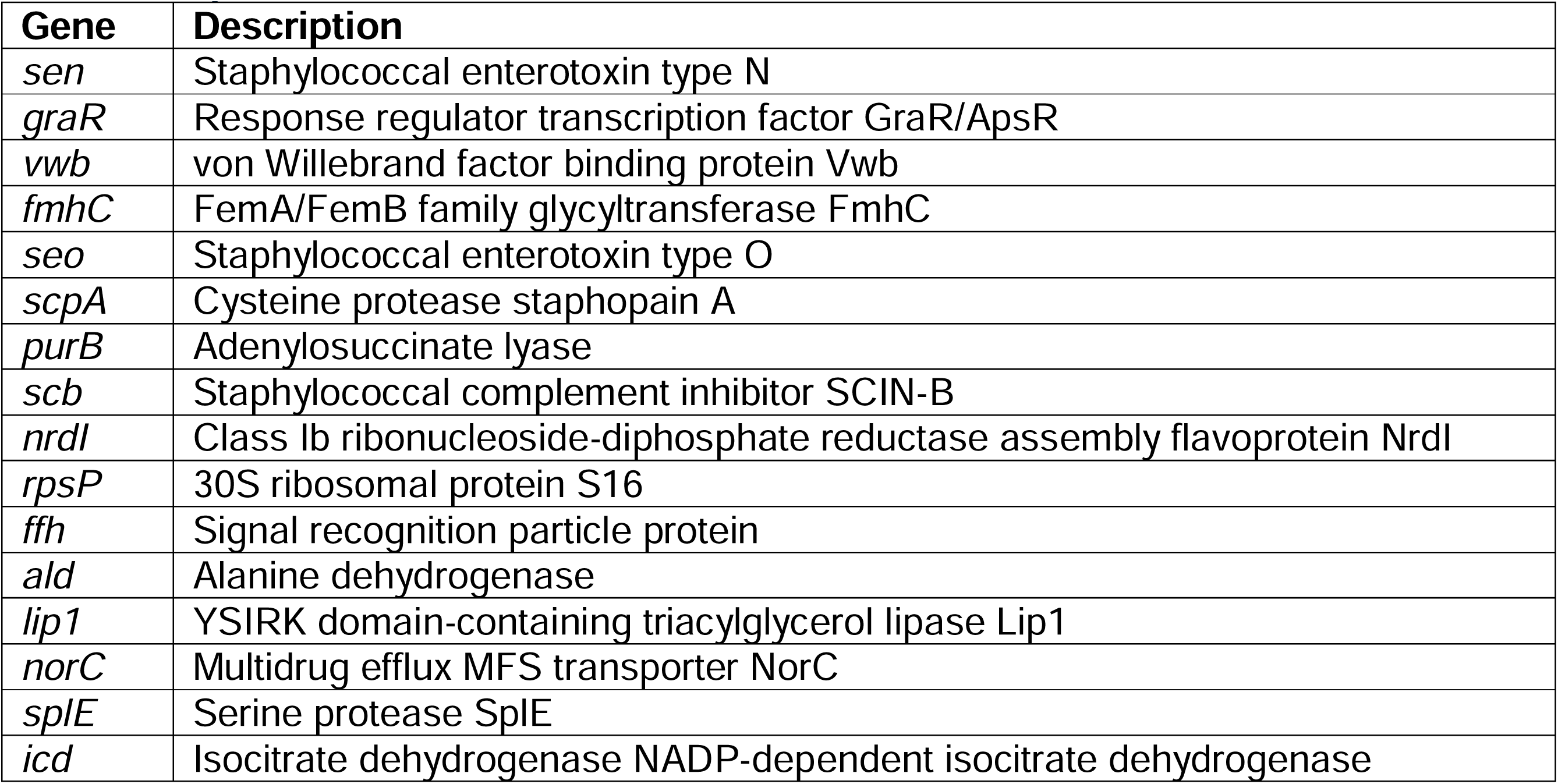
Genes containing k-mers significantly associated with the presence of SCC*mec* in *S. aureus* genomes (-log(p-value) > 200), ordered by descending maximum p-value.

K-mers that were significantly associated with SCC*mec* carriage, either more commonly present or more commonly absent, were mapped to a phylogeny of the *S. aureus* isolates (Figure 5). Fisher’s exact test was performed for each of the 38 k-mers to determine the nature of the correlation with SCC*mec* carriage (Supplementary Table 1). The majority of k-mers (87.5%, n = 35/40) exhibited positive correlations (OR >1), with 16 of the 35 showing highly significant covariation (p < 0.001). The only k-mer with a highly significant negative correlation (OR < 1, p = 0.0005) corresponded to *icd.* The higher prevalence of k-mers that are positively associated with SCCmec provides little evidence that there are elements that block SCC*mec* acquisition. Conversely, there is strong evidence that certain genes promote the integration or retention of SCC*mec.* This is consistent with possible potentiation of acquisition or compensatory change to accommodate SCC*mec* in the recipient genome.

**Figure 5:**
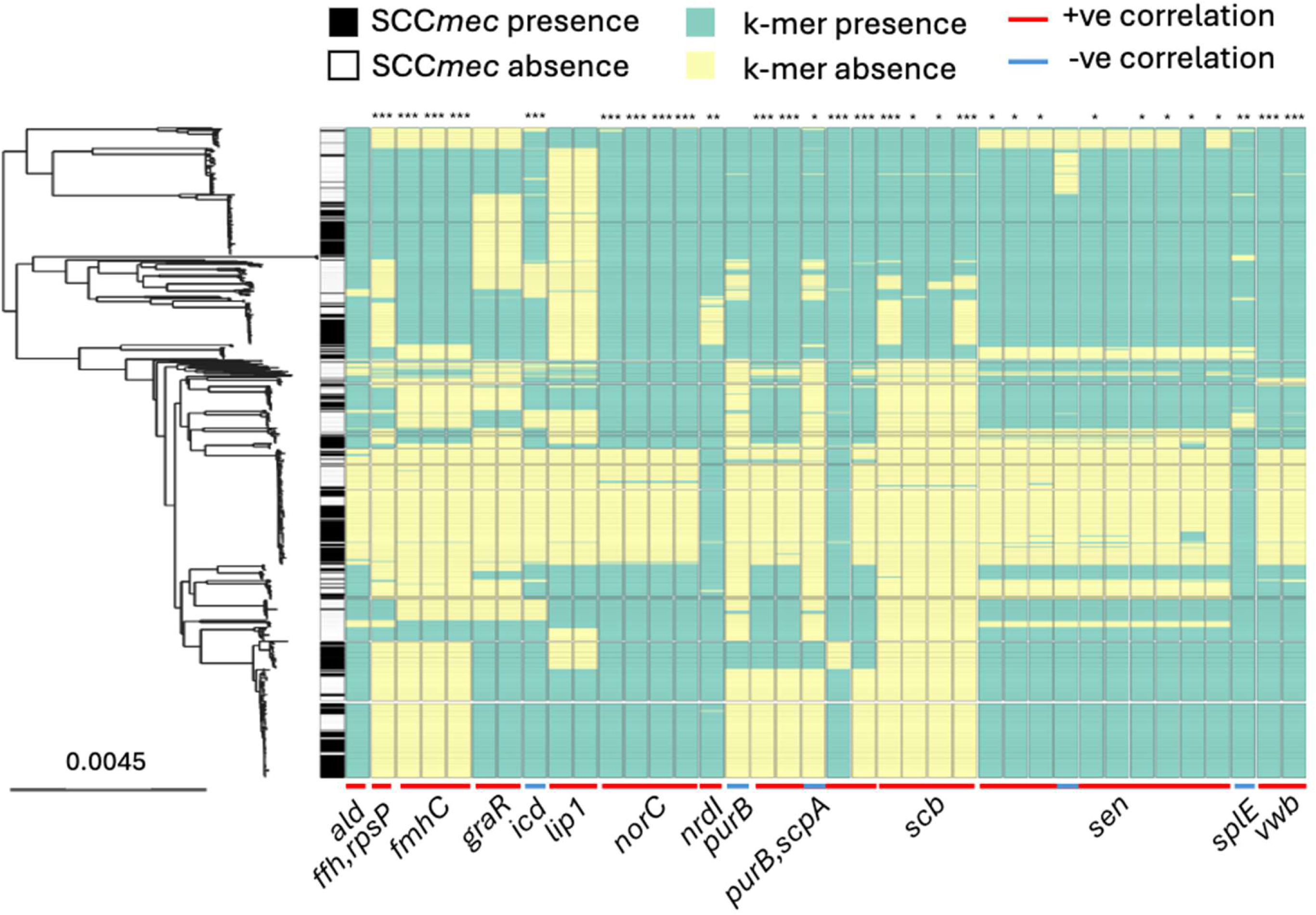
Presence of 38 k-mers significantly associated with SCC*mec* carriage. Maximum-likelihood phylogenetic heatmap showing the presence/absence of SCC*mec* (black/white) and 38 k-mers (green/yellow) that lie in coding regions and are significantly associated with SCC*mec* carriage. The corresponding genes are annotated below the heatmap. Created using Microreact (86). *: p < 0.05, **: p < 0.01, ***: p < 0.001.

## Discussion

There has been extensive work identifying genetic determinants of antimicrobial resistance in several bacterial species. While robust genotype-phenotype prediction is possible in some cases, accuracy is often less than 100% (50–53). Furthermore, where AMR is predicted from genome data, this is usually inferred for resistance above a certain threshold based on clinically relevant minimum inhibitory concentrations of antimicrobials used in laboratory bacterial growth assays. Of course, breaking phenotypes down into ‘yes / no’ metadata is an oversimplification. Even for relatively simple AMR phenotypes, where a single gene or allele may be the principal agent, there are complex genomic interactions that govern if expression occurs and to what extent (17,54–57).

Differentiating the nature of multiple gene interactions, such as synergistic additive effects and epistasis, is challenging from genome data alone. One of the main reasons for this is that the genomic signature of covariation is the same for coinheritance and LD, and functional co-adaption - here epistasis. Bench-marking the GOLD-GWAS approach with simulated data, we showed that it was possible to identify covarying alleles that were not explained by LD (Supp. Fig 1). Furthermore, extending the method to real data of *S. aureus* isolate genomes identified multiple significant associations among loci known to be functionally linked. In particular, *divIVA* and other genes involved in the peptidoglycan synthesis (*murC, yfhO, asp2, clfB, atl, bshB2*), including variants of genes encoding enzymes for uridine diphosphate N-acetylglucosamine (UDP-*GlcNAc*) biosynthesis (Table 1) (23,58–62).

Having demonstrated utility for detecting known adaptive covariation signatures, GOLD-GWAS was applied to identify genes that covary with SCC*mec* (Figure 3). Unsurprisingly, sequence variation at the known insertion site (*rlmH*) was strongly associated with SCC*mec* carriage, consistent with an established role in mobile genetic element integration and excision (25,49,63). Extending analyses with or without masking *rlmH,* detected k-mers that mapped to other genes, including some annotated as encoding hypothetical proteins (Figure 3B and Figure 4B). The most significant covariation with SCC*mec* was observed with the staphylococcal enterotoxin N gene (*sen*) (Figure 4B). This is associated with toxic shock-like syndromes and food poisoning (64) caused by staphylococci and epidemiological studies show that enterotoxin genes can be enriched in MRSA populations (65). This suggests a possible correlation between toxin production and antibiotic resistance, supported by the GOLD-GWAS analyses.

Among the other genes with sequence that co-varied with SCC*mec* were those linked directly linked to antibiotic resistance. The response regulator component of the glycopeptide resistance associated two-component system, *graR*, regulates susceptibility to vancomycin and daptomycin and is linked to cell wall stress responses (66–70). The relatively large average effect size reported here (beta = 0.8385) is likely attributable to the significant down-regulation of *mecA* in the absence of *graRS* (70). Furthermore, the *norC* gene encodes a multidrug efflux pump (71) and the von Willebrand factor binding protein (vWbp) have been linked to biofilm formation which improves survival rate under various stress conditions (72,73). These functions highlight the complex network of antibiotic resistance mechanisms that may interact with methicillin resistance conferred by *mecA*.

Quantifying and differentiating potential synergistic effects among virulence and resistance-associated loci is extremely important for understanding pathogen evolution. While direct evidence of epistasis requires phenotypic validation, we propose three adaptive scenarios that may explain putative SCC*mec* gene interactions identified in this study. First, specific loci could have a direct role in SCC*mec* insertion or excision by facilitating or hindering integration at the attB site. In this scenario, the presence or absence of specific regulatory or structural elements near the integration site may determine how efficiently SCC*mec* is inserted or excised from the bacterial chromosome. Second, some genes may undergo compensatory mutations following SCC*mec* insertion. In this case, mutations may balance the disruptive effects of integrating the large SCC*mec* element, restoring cell function and preserving bacterial fitness while enabling resistance (74–76). This scenario is consistent with the prevalence of positive correlations among k-mers that covary with SCC*mec*. Finally, there may be potentiation or functional synergy with methicillin resistance. Here, regulatory and virulence pathways, such as those govern by the GraRS two-component system, may interact with beta-lactam resistance, enhancing SCC*mec* effects by boosting resistance or altering the cell envelope to further reduce antibiotic susceptibility (77).

It is important to recognise the methodological limitations of GOLD-GWAS. Most importantly, computational genome analyses only provide statistical inference based upon sequence covariation. However, epistasis is measured phenotypically. Therefore, while our approach can identify candidates for further study, mechanistic understanding requires functional microbiological validation. There are also limitations of the analysis methodology. First, there is an emphasis on covariation with a target locus, rather than an all against all comparison. While this gives enhanced computational efficiency over some existing methods (42,78–81) and targeted gene identification, it inevitably overlooks multi-locus interactions that are not related to the gene(s) under investigation. Second, while masking LD regions helps to minimise confounding signals of coinheritance it may also mask potential functional interactions among genes in physical proximity.

## Conclusions

Integrated genome covariation analyses, such as that presented here, are an important step towards improved understanding of pathogen evolution. Context-free gene-centric approaches often fall short of accurate functional inference – such as AMR. With ever larger genome datasets and deeper understanding of gene function it is possible to move closer to accurate genotype-phenotype maps that account for gene network interactions.

## Methods

### Isolate genomes

All available *S. aureus* whole genome sequences were retrieved from the National Center for Biotechnology Institution (NCBI) reference sequence database with higher than 95% completeness or x30 coverage (n = 1,001, accessed 3rd October 2023). SCC*mec* regions were identified and typed using a custom database (github.com/Sheppard-Lab/sccmec_classifier) with minimap2 (v2.28) (82). Genomes with untypable SCC*mec* regions were removed from further analysis (n = 195). The final data set of 806 isolates consisted of 426 SCC*mec*-positive and 380 SCC*mec*-negative isolates. Isolate details including accession numbers are included in Supplementary Table 2. Core genome alignments were generated using PIRATE (v1.0.5) (83). A core genome maximum-likelihood phylogeny was created using RaxML (v8.2.12) (84) with GTRGAMMA as a substitution model. ClonalFrameML (v1.12) was used to account for recombination (85). All phylogenies were visualised using MicroReact (86)(87).

### Simulation data

Simulation data was generated using SimBac (44). Specifically, we simulated bacterial genomes with a recombination rate of R = 0.01 and a site-specific mutation rate of θ = 0.001 to generate 1,000 genomes, each spanning 1 Mbp. To introduce artificial genomic covariation, we identified a polymorphism located within 100 ±10 kbp with a minor allele frequency exceeding 0.2. Then, we generated eleven artificial covarying sites between 600,000 bp and 600,100 bp at 10 bp intervals. The nucleotide at each covarying site was determined based on the nucleotide present at the identified polymorphic site (at ∼100 kbp). The covariation was simulated with 95% accuracy to avoid perfect correlation, which could result in p-values of 0 and lead to numerical instability. The target region for masking was specified with start and end coordinates of 85896 bp and 101896 bp respectively.

### Computational masking

GOLD-GWAS is conceptually simple, involving: (i) estimating LD around a locus, here SCC*mec*; (ii) building a directory of unitigs and masking those within the LD region and the target region; (iii) conducting GWAS (Supplementary Figure 3). Linkage disequilibrium (LD) values were calculated using a custom pipeline (github.com/Sheppard-Lab/GOLD-GWAS/blob/main/run_ld_calculation.sh) Briefly, assemblies were aligned to the *S. aureus* NCTC 8325 reference genome using BWA (v0.7.17) (88). Alignments were then converted to BAM format, merged, sorted, and indexed with SAMtools (v1.16.1) (89). Variant-calling was performed by BCFtools (v1.14) (89). All single nucleotide polymophisms (SNPs) were counted, and 10% were randomly sampled to generate the variant call format (VCF) input file for LD analysis with PLINK (v1.9) (90). Only SNPs no more than 200,000 bps apart, using an LD window of 2,900 Mb for all SNPs, were analysed. LD decay plots were created with a custom R (v4.2.2) script (github.com/Sheppard-Lab/GOLD-GWAS/blob/main/plotting_lddecay. r).

The LD threshold was calculated as the intersection of the fitted R^2^ logarithmic bp decay and the average LD value, for distances exceeding 100 kbp, which are considered sufficiently large to represent a non-LD region (Figure 6A) (91). Due to potential regional variations in LD across the genome, a conservative cut-off of 10,000 bp was chosen for the masking procedure. The SCC*mec* target regions for masking were specified using the direct repeat and reverse complement inverse repeat (DR_SCC_-R and IR_SCC_-L, Supplementary Figure 1B) sequences as the start and end coordinates respectively, to capture all SCC*mec* types. From here, SCC*mec* and associated LD regions in SCC*mec*- positive isolates and LD regions in SCC*mec*-negative isolates were detected using the custom database option in ABRicate (v1.0.0)(92) with 80% coverage and 80% sequence identity thresholds (Figure 6B). Unitigs were generated from genome assemblies using the call mode of unitig-caller (v1.3.0) (93) with the pyseer flag and a k-mer size of 31. Unitigs were then mapped to the SCC*mec* (or *divIVA)* and associated LD region sequences using BWA (v0.7.17). Unitigs with over 80% sequence identity and 80% coverage were removed from further analysis (Figure 6C).

**Figure 6:**
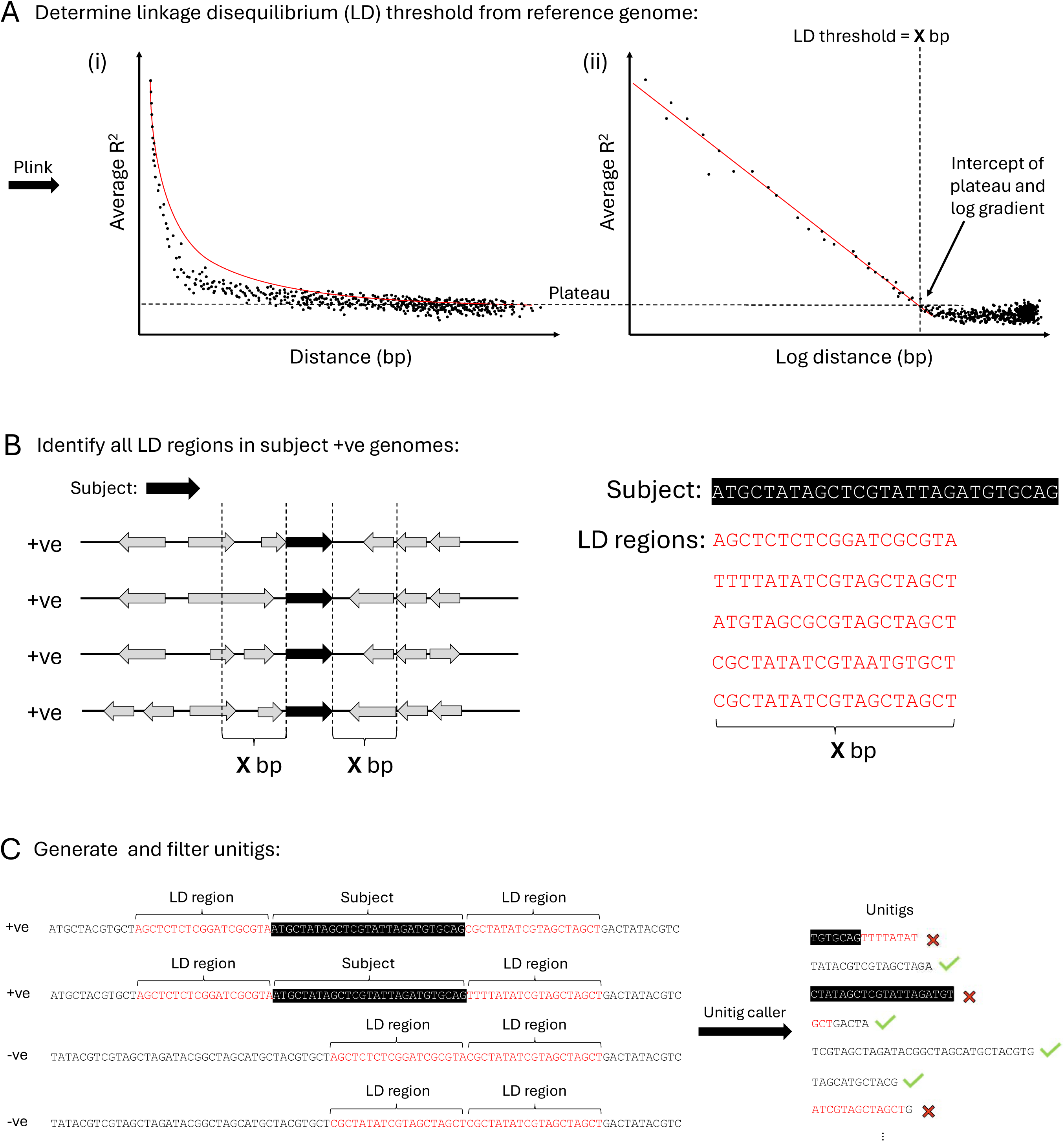
Computational Masking Method. (A) (i) Relationship between the linear distance between two SNPs and the average R^2^ value. Red line represents a polynomial trend line. (ii) Relationship between the logarithm of the distance between two SNPs and the average R^2^ value. Red line indicates the linear trend computed from the subset of samples excluding R^2^ < 0.2. The horizontal dotted line denotes the average R^2^ value observed beyond 100,000 bp, and the vertical dotted line marks the intersection of this with the trend line in the logarithmic distance plot. (B) Schematic demonstrating how sequences of LD regions are collected in association with the target gene based on the LD threshold from (A). (C) Unitigs with ≥80% identity with any target gene or LD region sequence are filtered out. Black boxes indicate target sequences, red text indicates the LD regions.

### Genome-wide association studies

GWAS was performed on the filtered set of unitigs using pyseer (v1.3.11) (94) with the linear mixed model flag. Manhattan plots were generated using Phandango (87) with JKD6159 (NCBI Reference Sequence: NC_017338.2) and HOU1444-VR (NCBI Reference Sequence: NZ_CP012593.1) as reference genomes for SCC*mec*-positive and SCC*mec*- negative strains, respectively. Gene hits were annotated using BWA (v0.7.17) with a minimum match length of 8 bp (88,94). The threshold for significance was calculated as

0.05 divided by the number of unique unitigs. From the hits that exceeded this threshold, the results were classified into two groups: those with -log(p-values) greater or less than the 3rd quantile (Q3) + 1.5x the interquartile range (IQR). To mitigate lineage effects (identified by poor chi-square values), we applied a minimum minor allele frequency threshold of 0.05%. To enhance biological relevance and reduce background noise, only genes with known names located in the vicinity of the identified k-mers were retained, while those with uncharacterised protein structures or functions were excluded from further analysis due to lack of information. To determine the nature of k-mer associations, Fisher’s exact test was performed using Python scipy.stats module fishers_exact with multiple testing correction implemented by statsmodel.stats.multitest. All scripts used for this analysis are available at github.com/Sheppard-Lab/GOLD-GWAS.

## Declarations

### Ethics approval and consent to participate

Not applicable.

### Availability of data and materials

All data analysed during the current study are included in this published article and its supplementary information files. Custom Python scripts used to perform analyses are available at github.com/Sheppard-Lab/GOLD-GWAS and zenodo (DOI: 10.5281/zenodo.15451691) unless otherwise stated in the text.

### Competing interests

The authors declare that they have no competing interests.

### Funding

SK is a self-funded PhD student and EAC is supported by a BBSRC grant (BB/W020602/1), awarded to SKS.

### Consent to publish

Not applicable.

### Authors’ contributions

SK: methodology, software, validation, formal analysis, writing. EAC: methodology, writing, visualisation. WM: conceptualisation, methodology. SKS: conceptualisation, writing.

## Supplementary

**Supplementary Figure 1:**
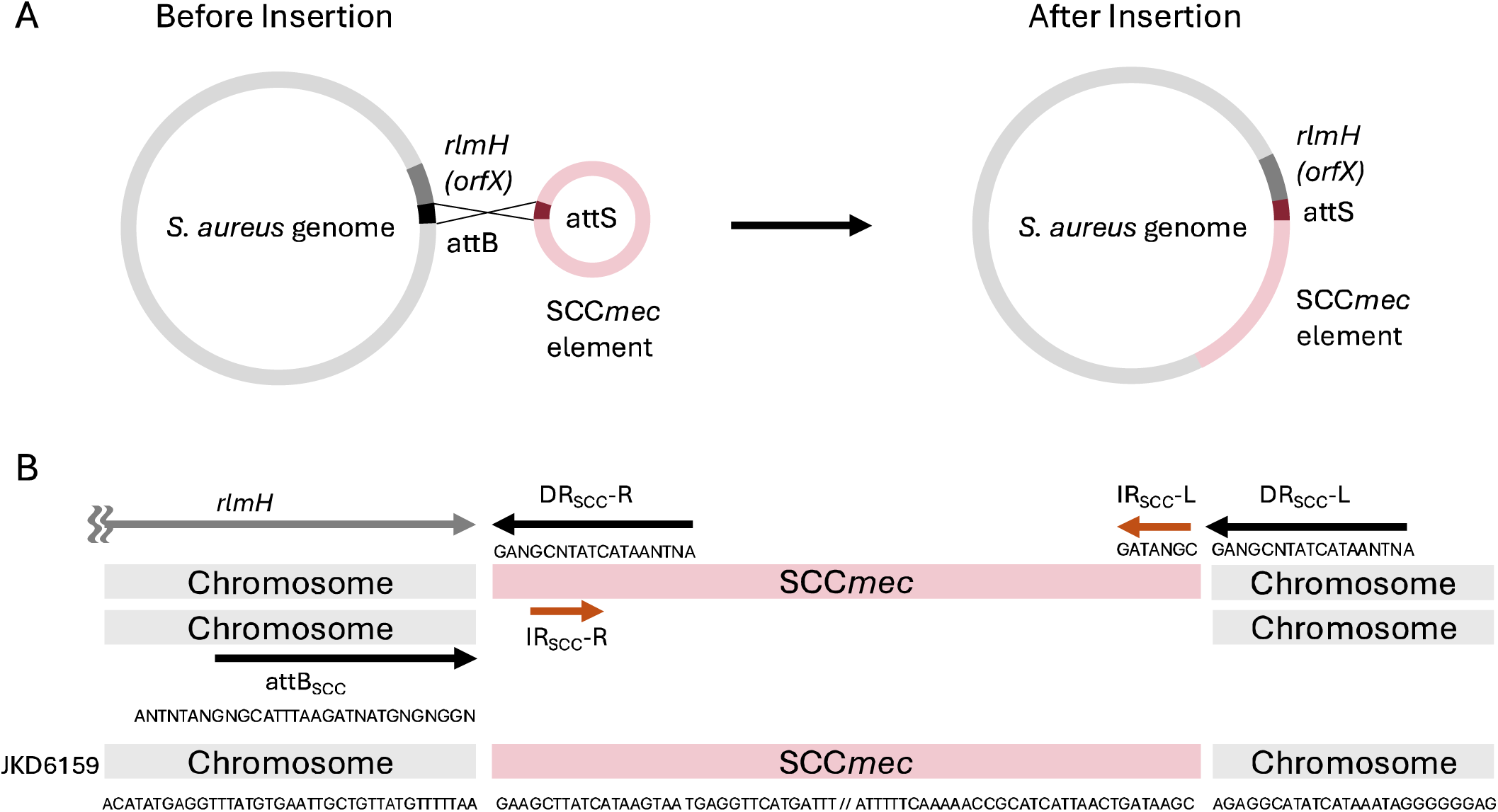
Site-specific integration of SCC*mec*. (A) The *S. aureus* genome before and after SCC*mec* insertion. The attB (blue) and attS (red) sites are responsible for SCC*mec* recombination. Before insertion, the attB site is located downstream of the *rlmH* sequence, which is replaced by attS once integration has occurred. (B) Structure of SCC*mec*. Within SCC*mec* there is one direct repeat (DR) sequence, and two reverse complement inverted repeats (IRs). An additional DR is present at the boundary of the downstream chromosomal region. Arrows indicate the orientation and positions of these repeats. Sequences based on the well-characterised clinical strain of *S. aureus* JKD6159 (NCBI Reference Sequence: NC_017338.2).

**Supplementary Figure 2:**
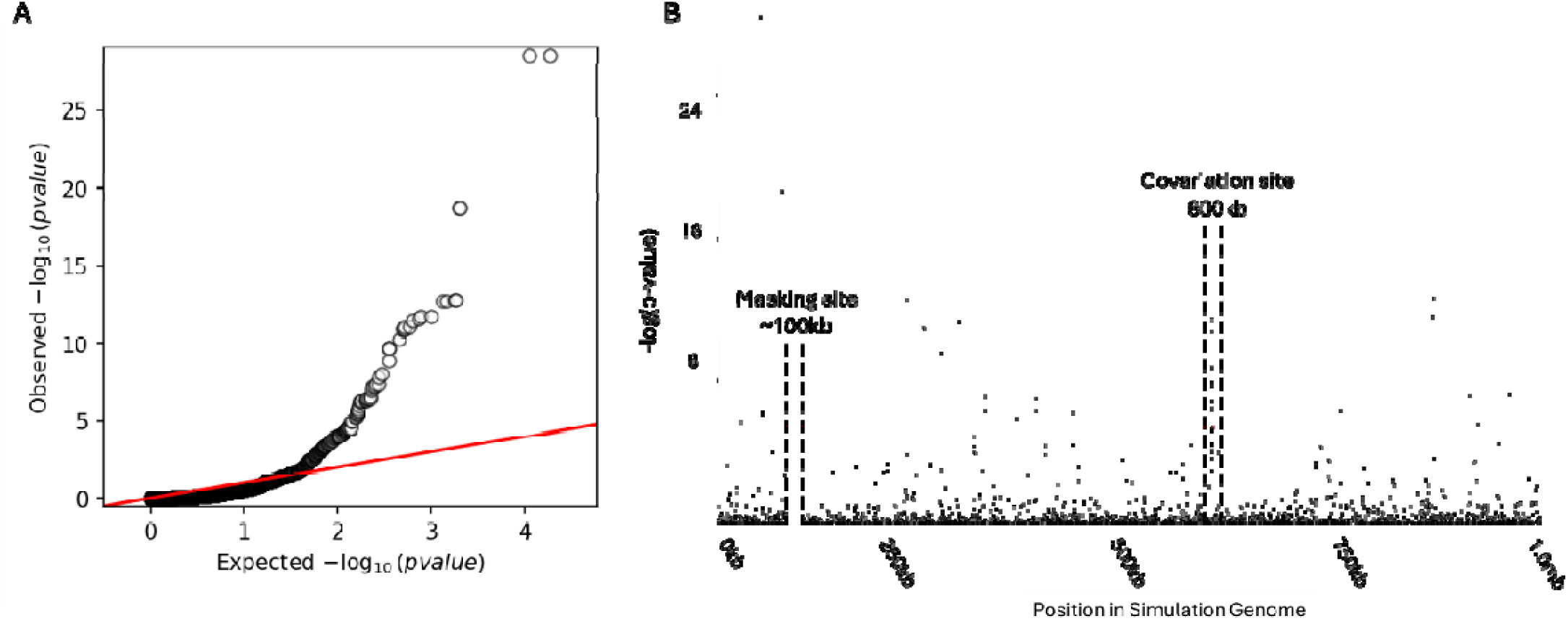
Summary of the genome-wide association study after masking an artificially created site of covariation. (A) Quantile-quantile plot comparing the expected -log(p-values) with observed -log(p-values). The red diagonal line indicates where the expected and observed values are equal. (B) Manhattan plot demonstrating the statistical significance association for selected variants arranged in order on a simulated genome. Each dot represents a k-mer. Red dotted line indicates the threshold for significance. Masked region near 100kb and artificial covariation region at 600kb are labelled.

**Supplementary Figure 3:**
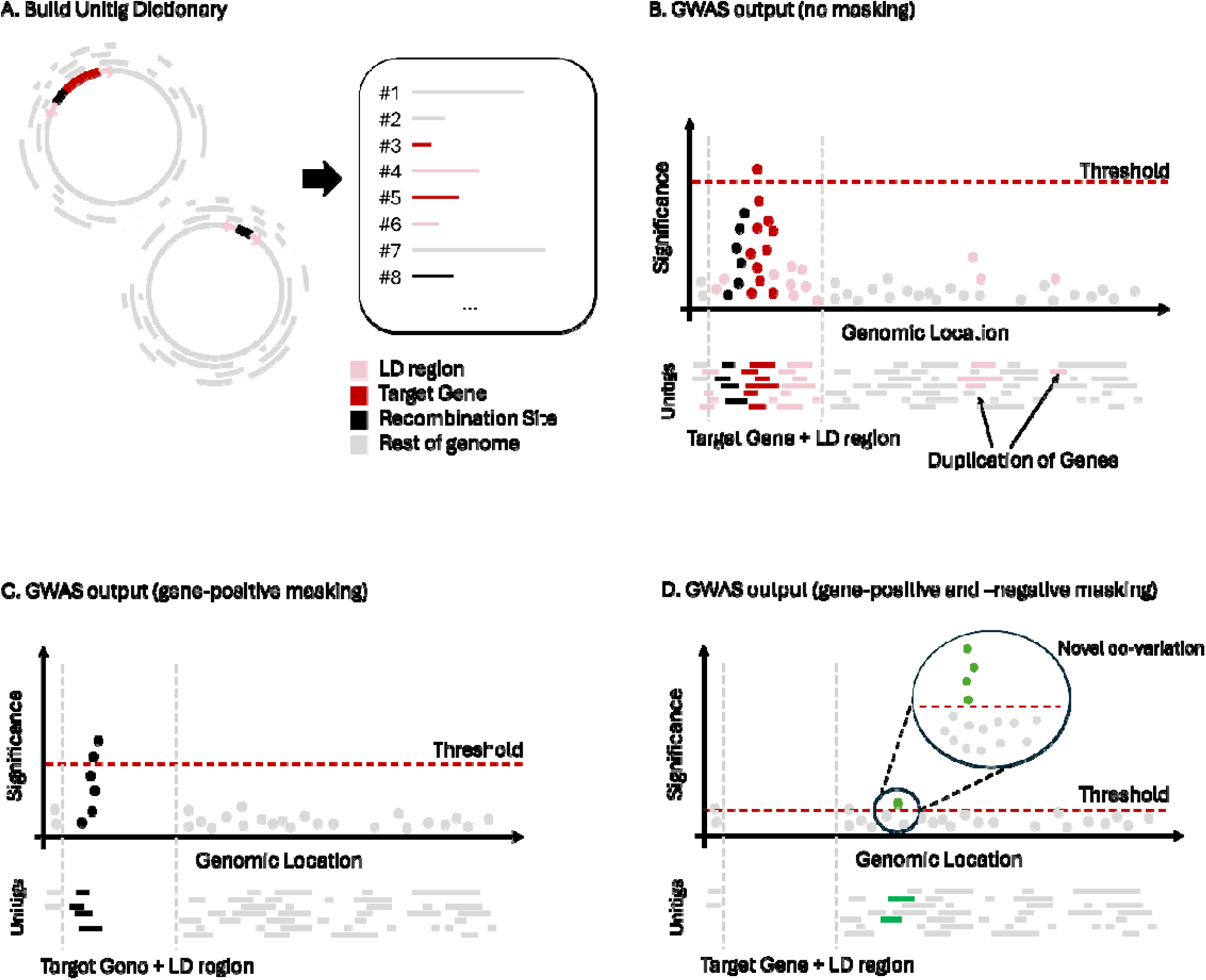
Summary of the GOLD-GWAS process. An overview conceptualising the methodology and advantages of GOLD-GWAS over standard GWAS. (A) A dictionary of unitigs is constructed that captures all genomic variation present in a pangenome. (B) Performing standard GWAS on this unitig dictionary using the presence of a target gene as the binary classifier will produce strong correlations for any unitigs that map to the target gene itself. Consequently, the extremely high p-values associated with these unitigs will result in a high threshold for statistical significance. Strongly associated unitigs may also include a recombination site, the target gene’s linkage disequilibrium regions, and any target gene unitig duplicated within the genome. (C) After identifying all target gene and LD region unitigs within gene-positive isolates, GOLD-GWAS removes these unitigs from the dictionary before performing GWAS. Unitig filtering now allows detection of recombination site or insertion sequence due to strong correlation with gene-negative isolates only. (D) By identifying all target gene and LD region unitigs in both gene-positive and -negative isolates and removing them prior to GWAS, GOLD-GWAS reveals previously hidden unitigs that significantly covary with the target gene.

## Notes

https://github.com/Sheppard-Lab/GOLD-GWAS

### Competing Interest Statement

The authors have declared no competing interest.

### Summary of Updates

Correction of typo in title; Figure numbers added to pdfs

