## Supplementary figures and images for "Genome co-adaptation and the evolution of methicillin resistant *Staphylococcus aureus* (MRSA)"

### Supplementary Figure 1

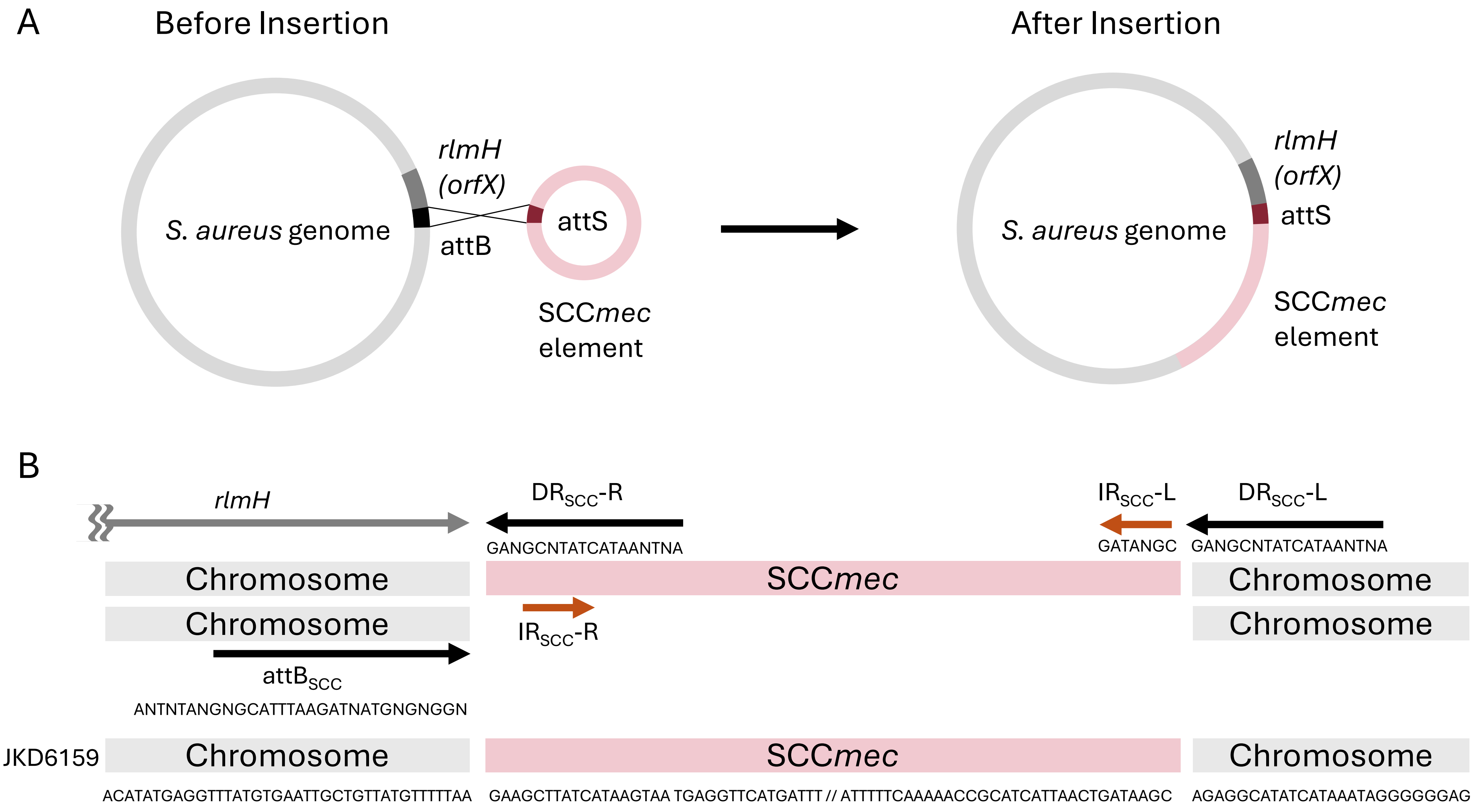

### Supplementary Figure 2

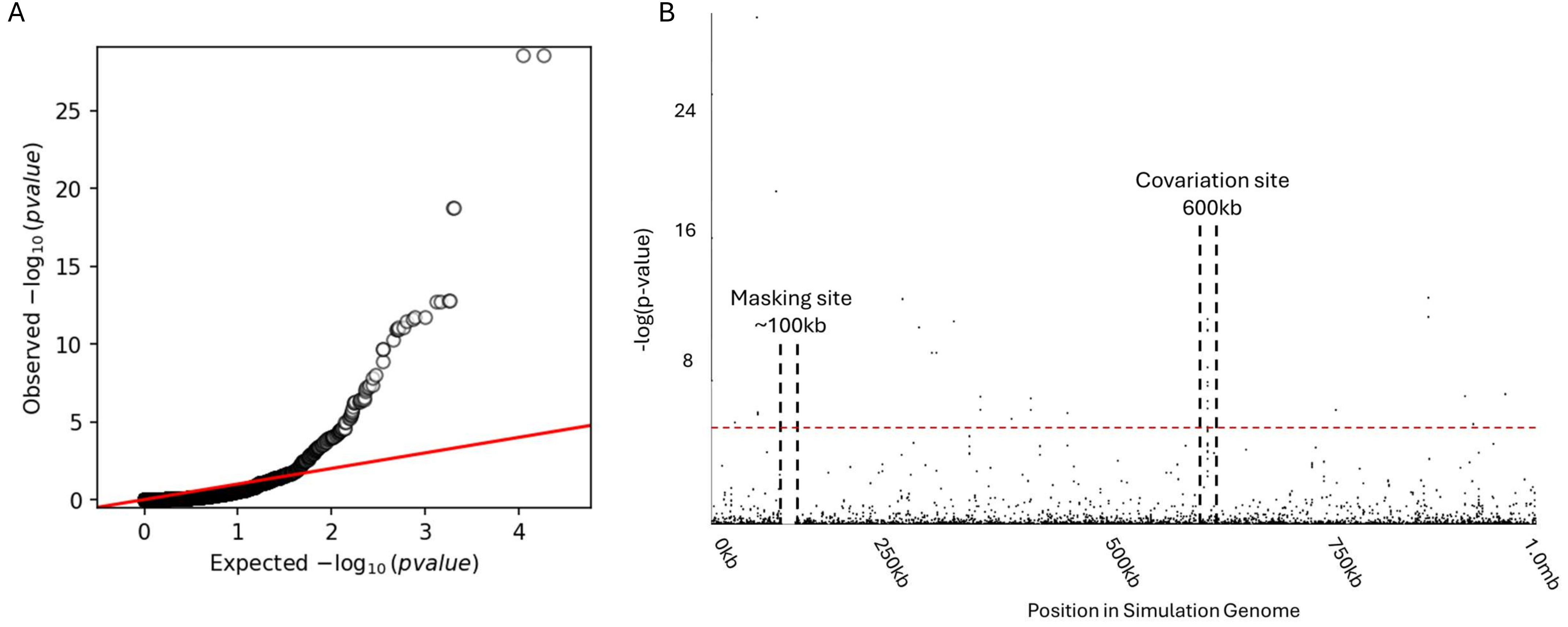

### Supplementary Figure 3

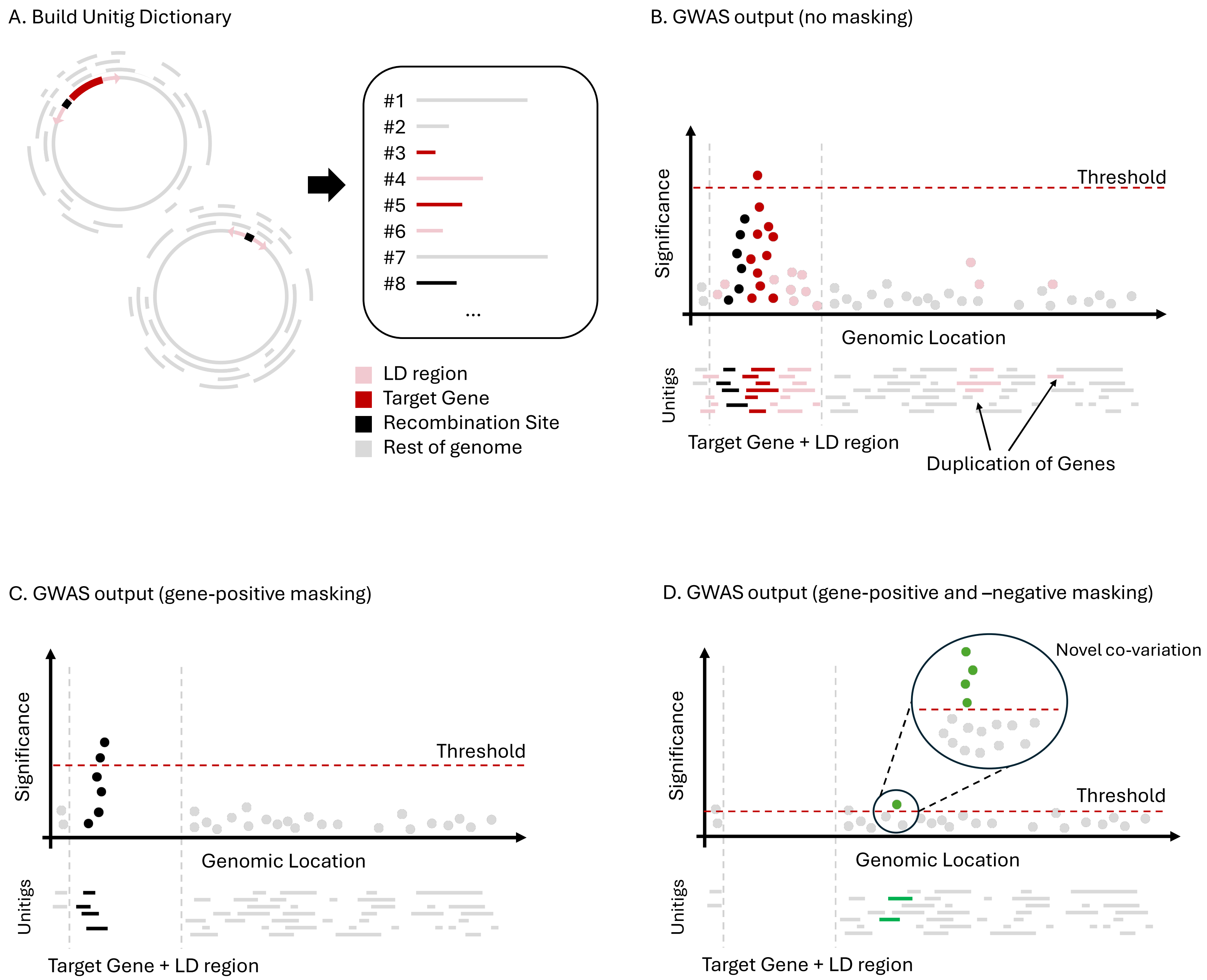
